# Perivascular signals alter global gene expression profile of glioblastoma and response to temozolomide in a gelatin hydrogel

**DOI:** 10.1101/273763

**Authors:** Mai T. Ngo, Brendan A.C. Harley

## Abstract

Glioblastoma (GBM) is the most common primary malignant brain tumor, with patients exhibiting poor survival (median survival time: 15 months). Difficulties in treating GBM include not only the inability to resect the diffusively-invading tumor cells but also therapeutic resistance. The perivascular niche (PVN) within the GBM tumor microenvironment contributes significantly to tumor cell invasion, cancer stem cell maintenance, and has been shown to protect tumor cells from radiation and chemotherapy. In this study, we examine how the inclusion of non-tumor cells in culture with tumor cells within a hydrogel impacts the overall gene expression profile of an *in vitro* artificial perivascular niche (PVN) comprised of endothelial and stromal cells directly cultured with GBM tumor cells within a methacrylamide-functionalized gelatin hydrogel. Using RNA-seq, we demonstrate that genes related to angiogenesis and remodeling are upregulated in the PVN model compared to hydrogels containing only tumor or perivascular niche cells, while downregulated genes are related to cell cycle and DNA damage repair. Signaling pathways and genes commonly implicated in GBM malignancy, such as *MGMT, EGFR*, PI3K-Akt signaling, and Ras/MAPK signaling are also upregulated in the PVN model. We describe the kinetics of gene expression within the PVN hydrogels over a course of 14 days, observing the patterns associated with tumor cell-mediated endothelial network co-option and regression. We finally examine the effect of temozolomide, a frontline chemotherapy used clinically against GBM, on the PVN culture. Notably, the PVN model is less responsive to TMZ compared to hydrogels containing only tumor cells. Overall, these results demonstrate that inclusion of cellular and matrix-associated elements of the PVN within an *in vitro* model of GBM allows for the development of gene expression patterns and therapeutic response relevant to GBM.

## 1. Introduction

Glioblastoma (GBM) is the most common primary malignant brain tumor. Prognosis is grim for GBM patients, with a median survival time of approximately 15 months [1]. Unlike many solid tumors where mortality is linked to metastasis, GBM tumors rarely metastasize [2, 3]. Instead, GBM spreads diffusively through the brain by invading along structural elements such as white matter tracts and blood vessels [3, 4]. Treatment is complicated by the diffuse spreading of tumor cells throughout the brain parenchyma. Poorly-defined tumor margins therefore limit the effectiveness of surgical resection, with tumor cells inevitably left behind after debulking. Subsequent radiation and chemotherapy have limited effect on the remaining tumor cells, with GBM tumors rapidly recurring (median: 6.9mo post debulking) at a site often in close proximity (>90% within 2cm) of the original resection cavity [1, 5]. The need for improved treatment methods for GBM highlights the importance of developing tools to facilitate the understanding of mechanisms of GBM invasion, the role played by interactions between tumor cells and the heterogeneous cell and matrix environment in the surrounding parenchyma, as well as to facilitate drug development and screening.

The heterogeneity of the GBM microenvironment presents a challenge in dissecting the biophysical, biochemical, and cellular cues that drive GBM invasion. Indeed, genomic and proteomic profiles of GBM that are derived from processing tumor tissue samples from patients or *in vivo* models can reflect contributions of both tumor and non-tumor cells that are present in the specimens [6, 7]. Therapeutic response as measured in *in vivo* models also reflects the synergistic effect of the drug on tumor and non-tumor cells. In particular, the microenvironment hosts a variety of cell types, including astrocytes, microglia, neural stem cells, pericytes, and endothelial cells, whose interactions with tumor cells have been demonstrated to contribute to heightened invasion potential [8]. Endothelial cells and stromal cells such as pericytes and fibroblasts are components of the perivascular niche, in which interactions between blood vessels and tumor cells have been implicated in a variety of roles that promote GBM progression [8, 9]. Tumor cells have been shown to associate closely with blood vessels through co-option as they invade into the surrounding brain parenchyma [4, 10, 11]. Co-opted vessels ultimately undergo regression, triggering hypoxic signaling (HIF-1) in the surrounding tumor tissue [12–16]. Subsequent secretion of pro-angiogenic factors such as VEGF induces sprouting angiogenesis at the tumor periphery [12, 15]. Tumor cells migrate towards these newly-formed blood vessels, and the cycle of co-option, regression, and angiogenesis repeats to push the invasive front of the tumor further into the brain parenchyma. Additionally, the perivascular niche potentially contributes to maintenance of cancer stem cells, a highly-resistant subpopulation of cells implicated in tumor recurrence [9, 17, 18]. Crosstalk between endothelial and cancer stem cells may be governed by Notch signaling, which promotes angiogenic activity and stem cell maintenance [8, 9, 17]. As such, further insight into the interactions between tumor and endothelial cells within the tumor core and tumor margins is critical towards developing strategies to inhibit the co-option, regression, and angiogenic mechanisms that contribute to tumor progression, as well as to potentially target the cancer stem cell subpopulation.

Investigating the interactions between multiple cell types traditionally utilizes conditioned medium in 2D cell culture or migration assays (e.g. Transwell ^®^). However, 2D culture has been shown to inaccurately predict therapeutic response, and direct cell-cell contact has been implicated in contributing to the interactions between heterogeneous cell types [19]. Notably, *Kebers et al*. reported that cell-cell contact was necessary to induce endothelial apoptosis by breast and fibrosarcoma cells, and *Borovski et al*. showed that proliferation of glioma propagating cells was sustained with direct contact with endothelial cells [19, 20]. Heterogeneous cell-cell contact is maintained in *in vivo* models, but the complexity of these platforms limits their utilization in determining how signaling interactions lead to observed phenotype. Thus, the drawbacks of 2D culture and *in vivo* models have paved the way for the development of 3D *in vitro* models, which utilize materials such as hydrogels to recapitulate elements of the tumor microenvironment [21–23]. To date, 3D *in vitro* models for GBM primarily incorporate only tumor cells, despite the demonstrated contributions of additional cell types to *in vivo* GBM behavior [24–31]. A few studies have examined the interactions between tumor cells, perivascular niche cells, and macrophages within 3D *in vitro* platforms, but these models feature spatial segregation between the different cell types that is not reminiscent of the *in vivo* microenvironment and prevents behavior that may result exclusively from direct cell-cell contact [32–35]. Mixed cultures of tumor cells, endothelial cells, and stromal cells have been achieved within biomaterials for breast, lung, and prostate cancers, and these models have been used to investigate how non-tumor cells impact tumor proliferation, phenotype, and therapeutic response [36–40]. Notably, physiologically-relevant behavior and therapeutic response have been observed in platforms that utilize model cell types for the non-tumor components, such as human umbilical vein endothelial cells (HUVECs) and mesenchymal stem cells (MSCs) or fibroblasts [37, 39–41]. Moreover, MSCs and fibroblasts have both been shown to exhibit pericyte-like behavior in supporting endothelial capillary formation [42].

Our lab has recently described a tri-culture biomaterial platform for the purpose of modeling the perivascular niche of GBM [43]. Previously, we demonstrated the formation of endothelial cell networks in gelatin hydrogels by co-culturing endothelial cells and fibroblasts. Using imaging, we observed that tri-culturing U87-MG tumor cells alongside these perivascular cells resulted initially in close association between U87-MG cells and developing endothelial cell networks, followed by dissolution of these networks in a manner similar to vessel co-option and regression *in vivo*. In this study, we trace the expression of select genes commonly implicated in GBM malignancy (e.g. *HIF1α, MMP9, TNC, EGFR*) and vessel co-option and regression (e.g. *ANG2* and *VEGF*) in correlation with the kinetics of endothelial network formation and dissolution observed in our platform. Additionally, we investigate how directly culturing perivascular niche cells alongside GBM tumor cells within a gelatin hydrogel alters the overall gene expression profile of the biomaterial, compared to hydrogels containing either perivascular niche cells or tumor cells. We use RNA sequencing (RNA-seq) to determine differentially-expressed genes resulting from tri-culturing perivascular cells with tumor cells, comparing results to cultures of tumor cells alone and perivascular cells cultured in the absence of tumor cells. We further investigate how the perivascular niche environment as a whole responds to temozolomide (TMZ), the frontline alkylating agent used clinically to treat GBM.

## 2. Materials and Methods

### 2.1. Cell Culture

Human umbilical vein endothelial cells (HUVECs) and normal human lung fibroblasts (NHLF) were purchased from Lonza (Walkersville, MD) and used before passage 6 and passage 8 respectively. HUVECs were cultured in EGM-2 media containing 2% FBS and supplements such as hEGF, hydrocortisone, VEGF, hFGF-B, R^3^-IGF-1, ascorbic acid, heparin, and gentamicin/amphotericin-B (Lonza, Walkersville, MD) [44]. NHLFs were cultured in FGM-2 media containing 2% FBS and supplements such as hFGF-B, insulin, and gentamicin/amphotericin-B (Lonza, Walkersville, MD) [45]. U87-MG cells, a gift from Professor Nathan D. Price (ISB, Seattle), were cultured in DMEM with 10% FBS and 1% penicillin/streptomycin [29]. All cell types were cultured at 37 °C in 5% CO_2_.

### 2.2. Fabrication of cell seeded gelatin hydrogels

Two distinct hydrogel formulations were used throughout all experiments. Both hydrogel variants were based on a methacrylamide-functionalized gelatin (GelMA) macromer previously developed in our lab for culture of GBM tumor cells [28] [29]. GelMA and HAMA were synthesized as previously described, and the degree of functionalization of GelMA was determined to be 50% by ^1^H NMR [29]. GelMA alone or GelMA combined with 15% w/w HAMA (GelMA + HA) were dissolved in PBS at 65 °C to form 4 wt% polymer solutions (0.6 wt% HAMA in GelMA + HA constructs). Lithium acylphosphinate (LAP) was added as a photoinitiator at 0.1% w/v for GelMA-only gels and 0.02% w/v for GelMA/HAMA gels. The reduced photoinitiator concentration allowed for equivalent elastic moduli (~3 kPa) between the GelMA and GelMA + HA constructs [43]. Cell seeded hydrogels (GelMA only; GelMA+HA) were then formed in one of three configurations: *GBM-only*; *Perivascular-only*; or *Tri-Culture. GBM-only* cultures were formed by re-suspending 1 × 10^5^ U87-MGs/mL in the GelMA or GelMA+HA polymer solution. *Perivascular-only* cultures were formed by re-suspending 1 × 10^6^ HUVECs/mL and 2 × 10^6^ NHLFs/mL in the GelMA or GelMA+HA polymer solution. *Tri-Cultures* were formed by re-suspending 1 × 10^6^ HUVECs/mL, 2 × 10^6^ NHLFs/mL, and 1 × 10^5^ U87-MGs/mL in the GelMA or GelMA+HA polymer solution. To visualize the interactions between the tumor and non-tumor cells, U87-MG cells were incubated with 10 μM CellTracker CMFDA (Thermo Fisher Scientific, Waltham, MA) prior to re-suspension in polymer solution. Cell-laden solutions were subsequently pipetted into circular Teflon molds with a diameter of 5 mm and a thickness of 1 mm, and exposed to UV light at a wavelength of 365 nm and intensity of 5.69 mW/cm^2^ for 30 s as previously described to form cell-seeded hydrogel disks [43]. Hydrogels were maintained for up to 14 days in 48-well plates in EGM-2 media with daily media changes.

### 2.3. Immunofluorescent staining and imaging

Formalin (Sigma Aldrich, St. Louis, MO) was used to fix *Perivascular-only* and *Tri-Culture* hydrogels. Cells were subsequently permeabilized with 0.5% Tween 20 (Thermo Fisher Scientific, Waltham, MA) in PBS, and blocked with 2% BSA (Sigma Aldrich, St. Louis, MO) and 0.1% Tween 20 in PBS. Hydrogels were stained using mouse anti-human CD31 (1:200, Dako, Denmark), rabbit anti-human ANG2 (1:100, Thermo Fisher Scientific, Waltham, MA), and/or goat anti-human VEGF (1:30, Novus Biologicals, Littleton, CO) as primary antibodies, and chicken anti-mouse Alexa Fluor 488 and donkey anti-rabbit Alexa Fluor 555 or donkey anti-goat Alexa Fluor 555 (1:500, Thermo Fisher Scientific, Waltham, MA) as secondary antibodies. Z-stacks were imaged using a DMi8 Yokogawa W1 spinning disk confocal microscope with a Hamamatsu EM-CCD digital camera (Leica Microsystems, Buffalo Grove, IL).

### 2.4. Flow cytometric analysis of Perivascular-only and Tri-culture hydrogels

*Perivascular-only* cultures (endothelial and stromal cells) and *Tri-Cultures* (GBM, endothelial, and stromal cells) were dissociated using a protocol adapted from *van Beijnum et al*. [46] at 3, 7, and 14 days post-culture. Briefly, cell-seeded hydrogels were dissociated in a solution of collagenase type IV (230 U/mL, Worthington, Lakewood, NJ), dispase (0.23 U/mL, Thermo Fisher Scientific, Waltham, MA), and hyaluronidase (8 U/mL, Sigma Aldrich, St. Louis, MO). The resulting cell solution was filtered through a 100 μm cell strainer (Thermo Fisher Scientific, Waltham, MA), centrifuged, and re-suspended in PBS with 0.1% sodium azide (Sigma Aldrich, St. Louis, MO), 1% penicillin/streptomycin, and 2% FBS. Cells were stained with Alexa Fluor 488-conjugated CD90 (5:100, R&D Systems, Minneapolis, MN; NHLFs) and propidium iodide (PI, 1:1000, Thermo Fisher Scientific, Waltham, MA; Dead cells). Flow cytometry was then performed using a BD LSR II flow cytometer (BD Biosciences, San Jose, CA). Data was subsequently analyzed using FCS Express 6.0 software (De Novo Software, Glendale, CA), first gating for cells (FSC-A/SSC-A), then viability through PI exclusion, then Alexa Fluor 488 fluorescence to determine the CD90+ fraction of each sample.

### 2.5. RNA Isolation and Quality Analysis

Hydrogels were collected at 0, 3, 7, and 14 days and stored at −80 °C until RNA extraction. RNA was extracted using an RNeasy Plant Mini Kit, followed by DNase digestion using a RNase-free DNase set (Qiagen, Hilden, Germany) [47]. RNA samples for RNA-seq were analyzed for quality using an Agilent 2100 bioanalyzer (Agilent, Santa Clara, CA). Samples with a minimum RIN of 7 were used for RNA-seq [48]. Quality testing of RNA was performed by the Roy J. Carver Biotechnology Center at the University of Illinois Urbana-Champaign.

### 2.6. RNA-seq analysis

RNA-seq was performed on RNA samples from the *Tri-Culture* hydrogels, as well as RNA samples comprised of a mixture of RNA from the *Perivascular-only* and *GBM-only* culture conditions (*Mixed*). Hydrogels were cultured for seven days. The comparison between *Tri-Culture* and *Mixed* samples was inspired by the methodology described by *Lilly et al*. [49]. *Mixed* samples were constructed such that the CD90+ fraction of the *Tri-Culture* and *Mixed* samples were equal. Libraries for RNA sequencing were prepared using the TruSeq Stranded mRNAseq Sample Prep Kit (Illumina, San Diego, CA), which fragments the mRNA, reverse transcribes it to cDNA, and adds sequence adaptors [50]. Libraries were subsequently quantitated with qPCR and sequenced by using the HiSeq 4000 sequencing kit version one in conjunction with a HiSeq 4000 (Illumina, San Diego, CA). 100 nt single-end reads were generated in 101 cycles to yield 55 – 70 million reads per sample. Library preparation and sequencing were performed by the Roy J. Carver Biotechnology Center at the University of Illinois Urbana-Champaign.

Library quality was checked using FastQC, and Trimmomatic [51] was used to remove low-quality reads. Trimmed libraries were subsequently aligned to the human genome (GRCh38.p10) using STAR [52], and featureCounts [53] was used to categorize the aligned reads to genes and other genomic features. 45393 genes were identified from the reads. Differential gene expression analysis was performed in R using limma and edgeR packages [54] [55]. Quality control was performed to filter out genes with few counts using a threshold of 0.5 counts per million, with 14269 genes remaining after applying the threshold. This accounted for >99% of the counts and 31% of the identified genes. TMM normalization was performed before determining differentially expressed genes. Functional annotation was performed, followed by gene ontology (GO) and KEGG pathway analyses using GOstats [56].

### 2.7. Real-time PCR

RNA was reverse transcribed to cDNA using reagents from the QuantiTect Reverse Transcription Kit (Qiagen, Hilden, Germany) and a Bio-Rad MyCycler thermal cycler (Bio-Rad, Hercules, CA). Real-time PCR was performed in an Applied Biosystems QuantStudio 7 Flex PCR System (Thermo Fisher Scientific, Waltham, MA) using Taqman Fast Advanced Master Mix and Taqman Gene Expression Assays (Thermo Fisher Scientific, Waltham, MA) (**Table 1**). RPS20 was used as the housekeeping gene [57]. Fold change was calculated using the ΔΔC_T_ method and normalized to expression at Day 0 in hydrogels without HAMA [58].

**Table 1.** Taqman Gene Expression Assays used for RT-PCR.

| Gene | Function | Taqman Gene Expression Assay ID |
| --- | --- | --- |
| RPS20 | Housekeeping | Hs03044413_gH |
| MMP9 | Extracellular matrix breakdown | Hs00957562_m1 |
| TNC | Extracellular matrix protein | Hs01115665_m1 |
| FLT1 | VEGF receptor; angiogenesis; vasculogenesis | Hs01052961_m1 |
| VEGFA | Endothelial migration and proliferation; angiogenesis | Hs00900055_m1 |
| EGFR | Receptor tyrosine kinase; proliferation; amplified in GBM | Hs01076090_m1 |
| PDGFB | Angiogenesis; vascular development; expressed in both GBM tumor and endothelial cells | Hs00966522_m1 |
| HAS2 | Hyaluronic acid synthesis | Hs00193435_m1 |
| PDGFRA | Receptor tyrosine kinase; expressed in GBM tumor cells | Hs00998018_m1 |
| TEK | Binds to ANGPT1 and ANGPT2; involved in vascular development | Hs00945150_m1 |
| PDGFRB | Receptor tyrosine kinase; expressed in GBM endothelial cells and pericytes | Hs01019589_m1 |
| HIF1A | Regulates response to hypoxia; angiogenesis | Hs00153153_m1 |
| RHAMM | Receptor for hyaluronic acid; cell motility | Hs00234864_m1 |
| HAS1 | Hyaluronic acid synthesis | Hs00758053_m1 |
| ANGPT1 | Angiogenesis; vascular remodeling and stability | Hs00919202_m1 |
| ANGPT2 | Angiogenesis; expressed by co-opted vessels | Hs00169867_m1 |

### 2.8. Analysis of culture response to temozolomide (TMZ)

*Tri-culture* and *GBM-only* hydrogels were developed with and without HAMA and cultured for 7 days in EGM-2 media with daily media change. After 7 days of culture, TMZ (Sigma Aldrich, St. Louis, MO) was added at 0, 100, 250, and 500 μM to all hydrogels for 48 hours. To assess cellular response to TMZ, metabolic activity was assessed using a 3-(4,5-dimethylthiazol-2-yl)-2,5-diphenyltetrazolium bromide (MTT) assay (Thermo Fisher Scientific, Waltham, MA) 48 hours after treatment [59]. Hydrogels were incubated in 200 μL EGM-2 media and 20 μL MTT solution for 4 hours at 37 °C. The resultant formazan was solubilized into 300 μL DMSO (Sigma Aldrich, St. Louis, MO) overnight. Absorbance was read at 540 nm using a BioTek Synergy HT microplate reader (BioTek, Winooski, VT). Metabolic activity for treated samples was normalized to the untreated control. Additionally, growth rate inhibition (GR), as described in *Hafner et al*. [60], was measured in hydrogels at concentrations of 0, 25, 100, 250, and 500 μM TMZ after 7 days of culture. Relative cell count was assessed after 48 hours after treatment with the CellTiter-Glo Luminescent Cell Viability assay (Promega, Madison, WI) using a protocol derived from *Lin et al*. [61]. Briefly, hydrogels were incubated for one hour at room temperature in 150 μL EGM-2 media mixed with 150 μL CellTiter reagent. Luminescence was read using a BioTek Synergy HT microplate reader. Initial cell counts were obtained by running the assay on untreated hydrogels on the day of treatment. GR metrics were calculated as described by *Hafner et al*. [60], and the GR_50_ was calculated using the Online GR Calculator (www.grcalculator.org/grcalculator). DMSO at the highest relative concentration was used as the untreated control (0 μM).

### 2.9. Statistics

Statistical analysis for real-time PCR was determined using Kruskal-Wallis ANOVA followed by Dunn’s test for comparisons across time points, and Mann-Whitney test for comparisons with and without HAMA at a given time point in OriginPro (OriginLab, Northampton, MA). Differential gene expression, gene ontology, and KEGG pathway analyses were performed in R as previously described (*2.6. RNA-seq analysis*). Statistical analysis for drug treatment experiments was performed using either student’s t test or Mann-Whitney test for comparisons at a given drug concentration. Flow cytometry analysis and RNA-seq were determined using n = 3 hydrogels per group. Real-time PCR, metabolic activity, and growth rate inhibition results were determined using n = 6 hydrogels from two independent experiments. Significance was set at p < 0.05 for all analyses except for differential gene expression, in which an FDR adjusted p-value < 0.05 was used. Data are presented as mean ± SD.

## 3. Results

### 3.1. Gene expression changes accompany process of endothelial network co-option and regression

We cultured *Perivascular-only* and *Tri-culture* hydrogels for up to fourteen days (**Figure 1a**). We observed association between tumor cells and developing endothelial networks by three days of culture (**Figure 1b**). Endothelial networks continued to develop over fourteen days of culture in *Perivascular-only* hydrogels, but regressed by Day 14 in *Tri-culture* hydrogels (**Figure 1c, Supplementary Figure 1**). We performed staining for ANG2 and VEGF to investigate their distribution amongst CD31+ endothelial cells (**Supplementary Figures 2-3**). ANG2 was expressed in endothelial cells, and in particular was present in developed endothelial network structures at Day 7 and regressed cells at Day 14. VEGF was concentrated in and around developing endothelial network structures at Day 3, and became more diffuse with time.

**Figure 1.**
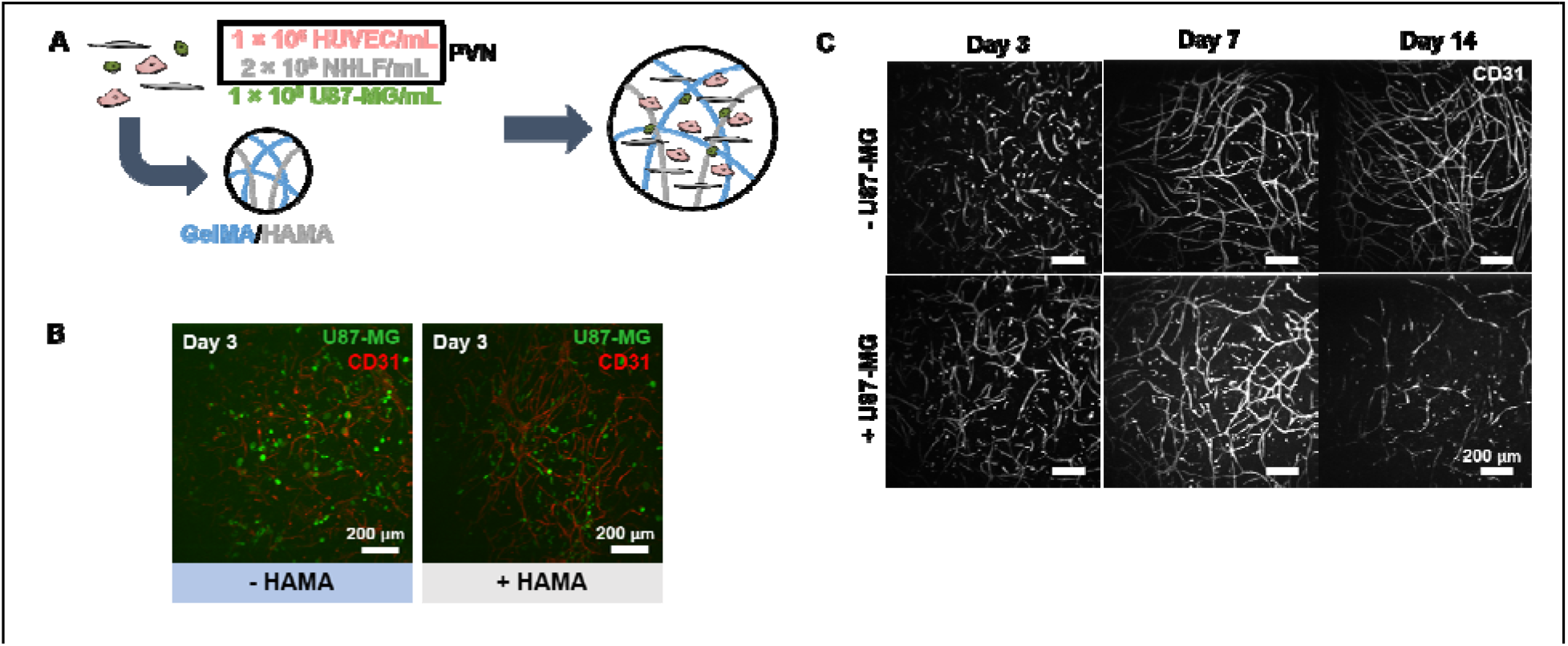
**(A)** Co-culturing perivascular niche (PVN) cells and U87-MG tumor cells inside gelatin hydrogels to develop an artificial GBM PVN. **(B)** CellTracker-labeled U87-MG tumor cells are located in close proximity to CD31+ endothelial cell networks after three days of culture. Scale bar: 200 μm. **(C)** Dissolution of endothelial networks is dependent upon presence of U87-MG cells in culture. Images are taken of GelMA-only constructs. Scale bar: 200 μm.

We subsequently used real-time PCR to investigate the kinetics of gene expression in *Tri-Culture* hydrogels during the process of endothelial network regression (**Figures 2 - 3**). We examined a panel of genes related to receptor tyrosine kinases (RTKs) (e.g. *EGFR, VEGFR1, TIE2, PDGFRA, PDGFRB*), hyaluronic acid synthesis and interactions (e.g. *RHAMM, HAS1, HAS2*), angiogenesis (e.g. *ANG1, ANG2, VEGF, HIF1α, PDGFB*), and ECM remodeling (e.g. *MMP9, TNC*) known to be associated with vessel maturation as well as co-option and regression. In GelMA hydrogels, we observed statistically significant decreases in all RTKs except for *EGFR* and *PDGFRα* with time (**Figure 2a**); however, we note that *PDGFRα* expression trended downwards with time. Within genes related to hyaluronic acid, we observed that *HAS1* expression trended upwards while *HAS2* expression decreased with time (**Figure 2b**). With regards to angiogenesis, we observed statistically significant decreases in *ANG2* and *PDGFB* expression with time, while *HIF1α* remained constant and *VEGF* trended upwards (**Figure 2c**). Finally, we observed a statistically significant increase in *MMP9* expression with time, as well as upregulation of *TNC* compared to Day 0 (**Figure 2d**).

**Figure 2.**
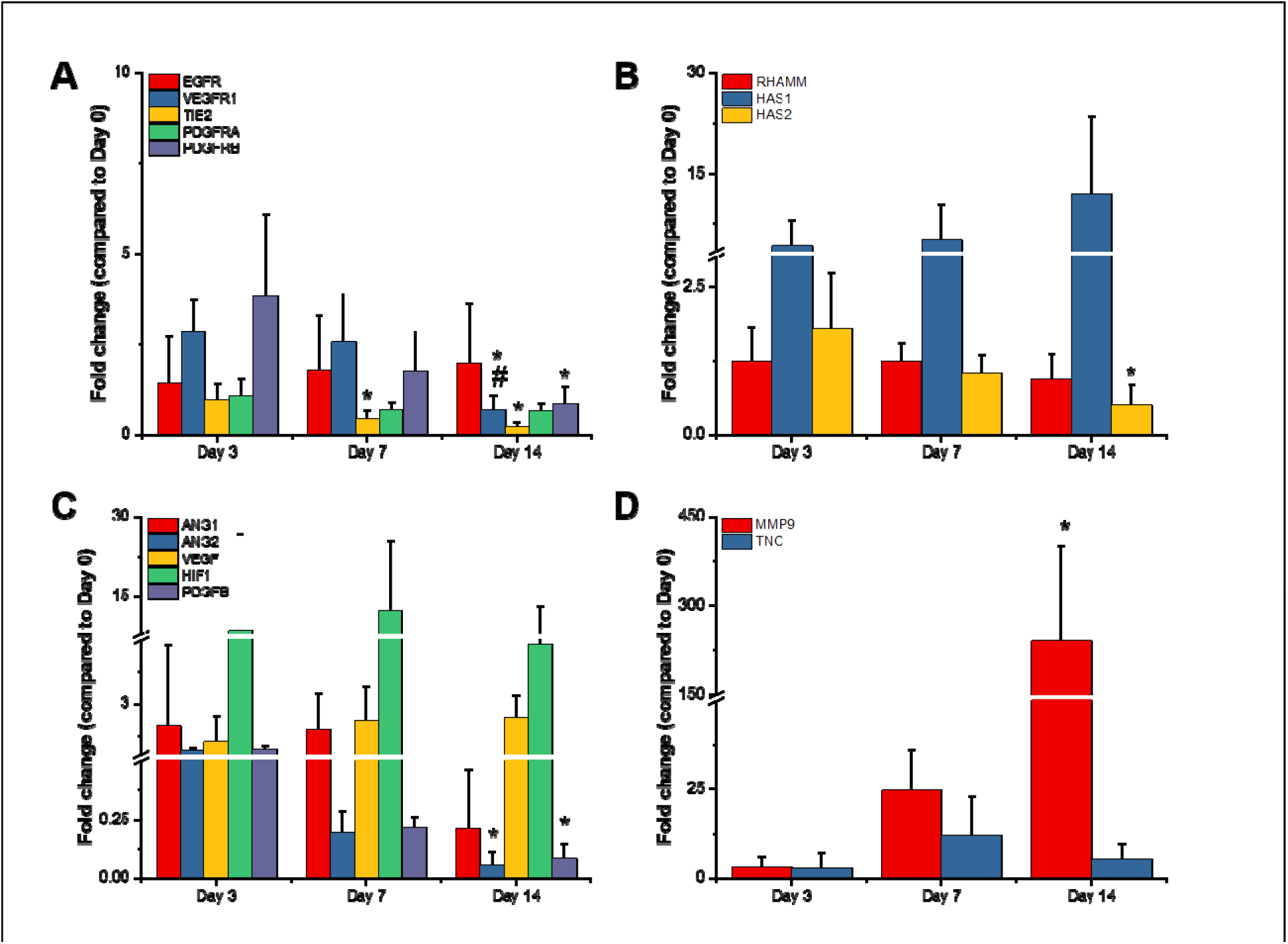
Gene expression profiles for GelMA-only *Tri-Culture* hydrogels over the course of 14 days. Genes studied were related to **(A)** receptor tyrosine kinases, **(B)** hyaluronic acid synthesis and interactions, **(C)** angiogenesis, and **(D)** ECM remodeling. *p<0.05 compared to Day 3. #p<0.05 compared to Day 7. Data is presented as mean ± SD. Two independent experiments were performed with three replicates each.

**Figure 3.**
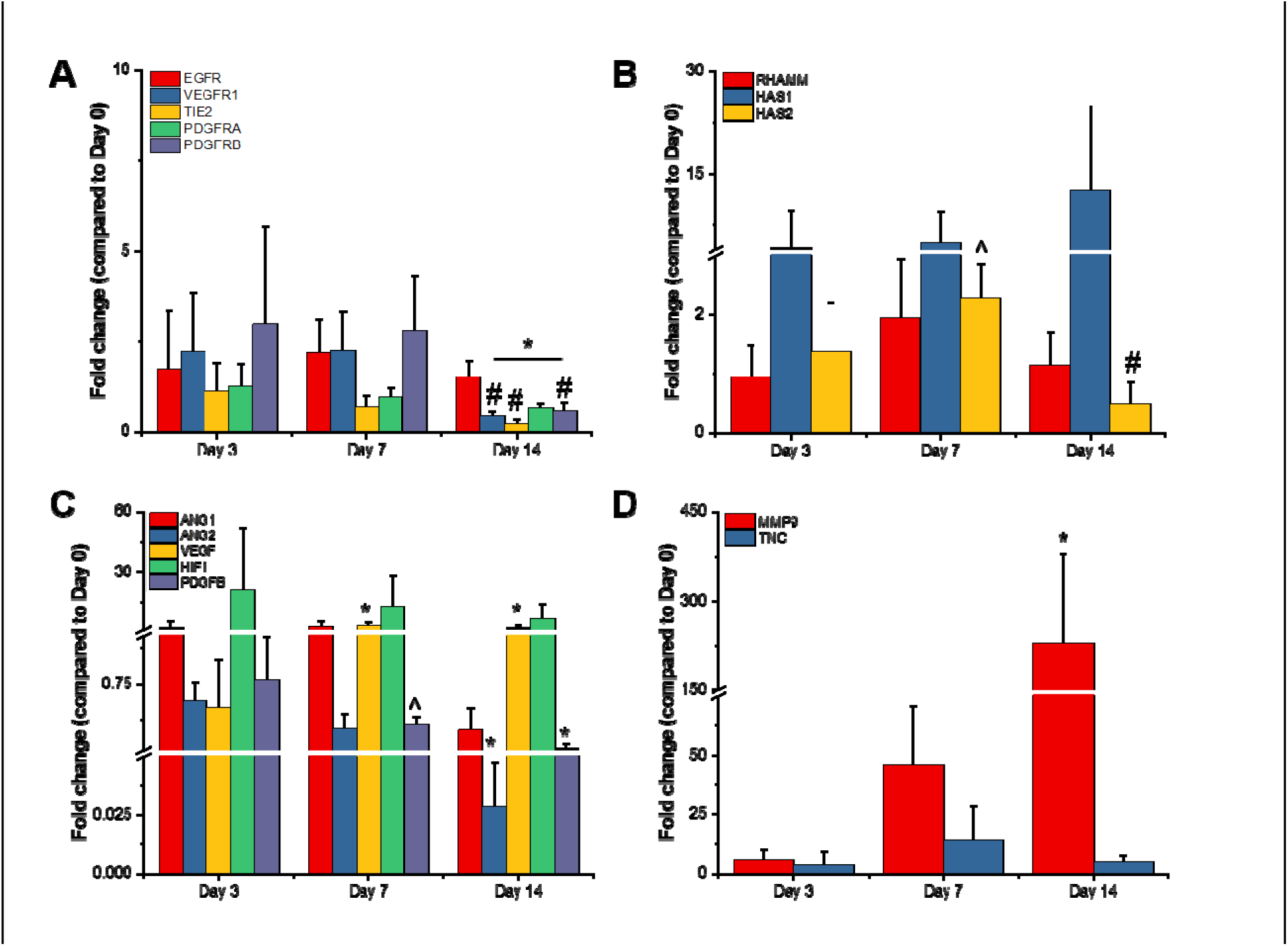
Gene expression profiles for GelMA + HAMA *Tri-Culture* hydrogels over the course of 14 days. Genes studied were related to **(A)** receptor tyrosine kinases, **(B)** hyaluronic acid synthesis and interactions, **(C)** angiogenesis, and **(D)** ECM remodeling. *p<0.05 compared to Day 3. #p<0.05 compared to Day 7. ^^^p<0.05 compared to GelMA-only hydrogel at same time point. Data is presented as mean ± SD. Two independent experiments were performed with three replicates each.

### 3.2. Gene expression kinetics are mostly conserved in the presence of covalently-bound hyaluronic acid

We observed that gene expression patterns for *Tri-Cultures* in GelMA + HAMA hydrogels mostly paralleled those observed in GelMA hydrogels (**Figure 3**). Statistically significant differences due to the presence of HAMA were only observed at Day 7 for *HAS2* AND *PDGFB* expression (**Figure 3b-c**). In particular, *HAS2* expression appeared to peak at Day 7, while expression continually trended downwards in GelMA hydrogels. While not statistically significant, we also observed that *HIF1α* expression trended downwards with time with the addition of HAMA, while expression was relatively constant in GelMA hydrogels.

### 3.3. Culturing perivascular niche cells along with GBM tumor cells alters overall gene expression profile of gelatin hydrogel

We subsequently used RNA-seq to analyze the gene expression profile of *Tri-Culture* GelMA-only hydrogels in contrast with *Perivascular-only* and *GBM-only* GelMA-only hydrogels cultured for seven days. RNA from the *Tri-Culture* hydrogels (*Tri-Culture*) was compared to RNA samples (*Mixed*) constructed from combining RNA from separate *Perivascular-only* and *GBM-only* hydrogels (**Figure 4**), such that the CD90+ fraction (NHLF sub-population) of the mixed samples would match that in the *Tri-Culture* samples (**Table 2**). Constructing RNA samples with similar cellular compositions allowed us to minimize the effect of disparate cell populations on differential gene expression results, thereby enabling us to identify the effect of culturing tumor and perivascular niche cells in direct contact in the same biomaterial, compared to segregated cultures of these cell types. We confirmed that CD90, along with the common endothelial marker CD31, were not differentially expressed between the *Tri-Culture* and *Mixed* samples (not shown). Differential gene expression analysis identified 1737 statistically significant upregulated genes (FDR-adjusted p < 0.05) and 1778 statistically significant downregulated genes (FDR-adjusted p < 0.05) in the *Tri-Culture* compared to the *Mixed* samples (**Figure 5, Supplementary Tables 1-2**). Samples clustered together based on whether they were *Tri-Culture* or *Mixed* samples (**Figure 5**). Gene Ontology (GO) analysis revealed that upregulated genes due to culturing GBM cells in direct contact with perivascular cells were correlated to angiogenesis and vasculature development; signaling and stimuli response; ECM remodeling, organization, and disassembly; and motility and adhesion (**Table 3, Supplementary Table 3**). Additionally, we noted upregulation of several fibrillar matrix components (e.g. *CSPG4, FN1, TNC*). KEGG pathway analysis additionally demonstrated over-representation of signaling pathways related to cell survival, growth, and migration amongst upregulated genes (e.g. PI3K-Akt, Ras, MAPK, Jak-Stat) (**Figure 6, Supplementary Table 5**). Notably, several receptor tyrosine kinases (RTKs) that feed into these pathways were upregulated, such as *EGFR, PDGFR*, and *FGFR1*. Moreover, HIF-1 and Notch pathways were over-represented, as well as ECM-receptor interactions. With regards to therapeutic response, we noted that *MGMT* was upregulated, and the KEGG term “EGFR tyrosine kinase inhibitor resistance” was over-represented amongst upregulated genes in response to *Tri-Culture* of GBM cells with perivascular networks. This was coupled with downregulated mismatch repair pathway genes. Both GO and KEGG analyses revealed that downregulated genes due to *Tri-Culture* of GBM cells with perivascular networks were broadly correlated to cell cycle and DNA replication; metabolism; and DNA damage repair (**Table 4, Supplementary Tables 4,6**).

**Figure 4.**
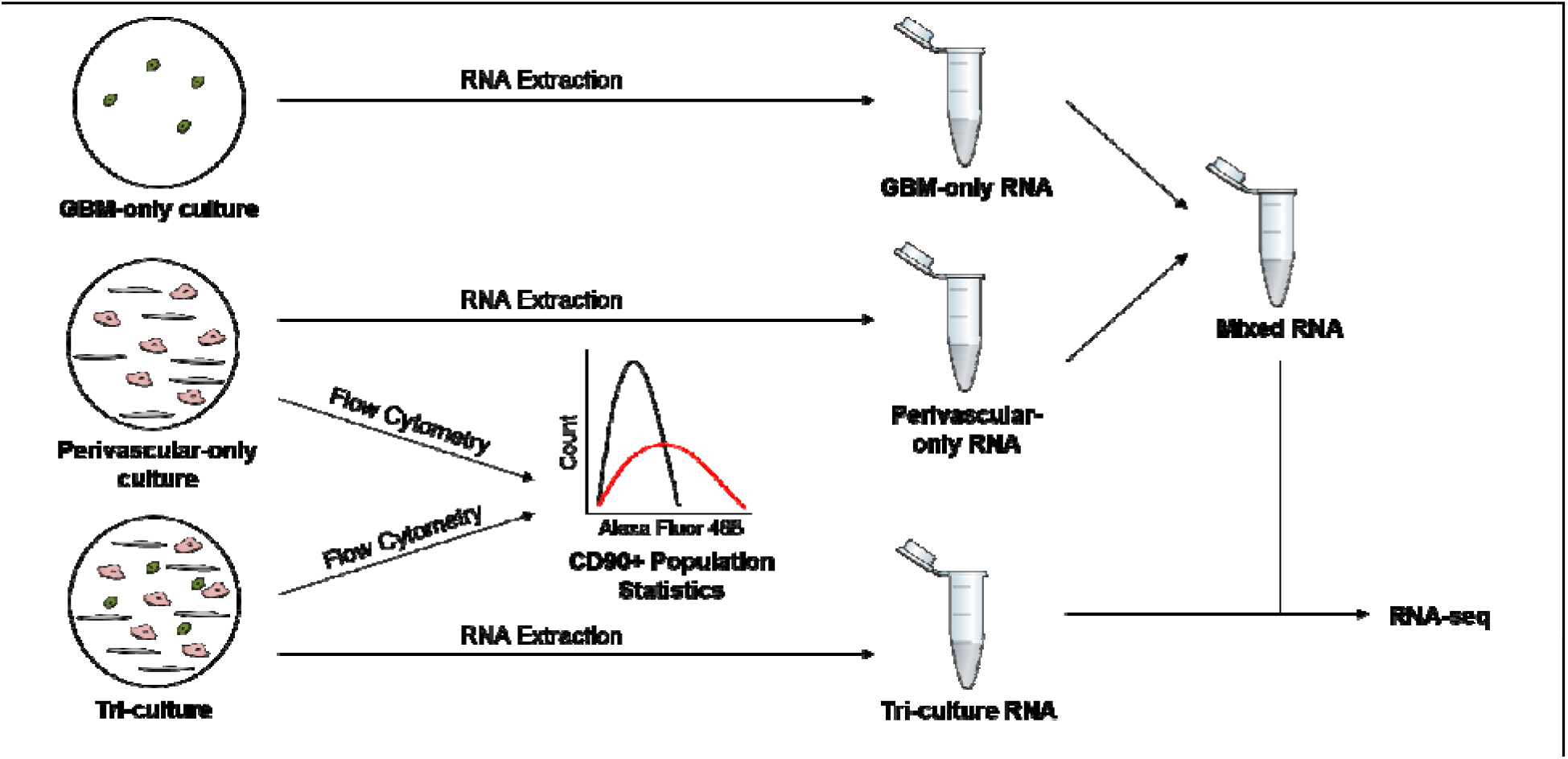
Schematic of workflow for preparing RNA samples for RNA-seq. RNA was extracted from *GBM-only, Perivascular-only*, and *Tri-Culture* hydrogels. Flow cytometry was performed on cells from additional *Perivascular-only* and *Tri-Culture* hydrogels to obtain the fraction of CD90+ cells in each culture system. *GBM-only* and *Perivascular-only* RNA were subsequently combined to form a *Mixed* sample such that the CD90+ fraction of the *Mixed* sample was equal to that in the *Tri-Culture* sample. *Mixed* and *Tri-Culture* RNA were subsequently sequenced.

**Figure 5.**
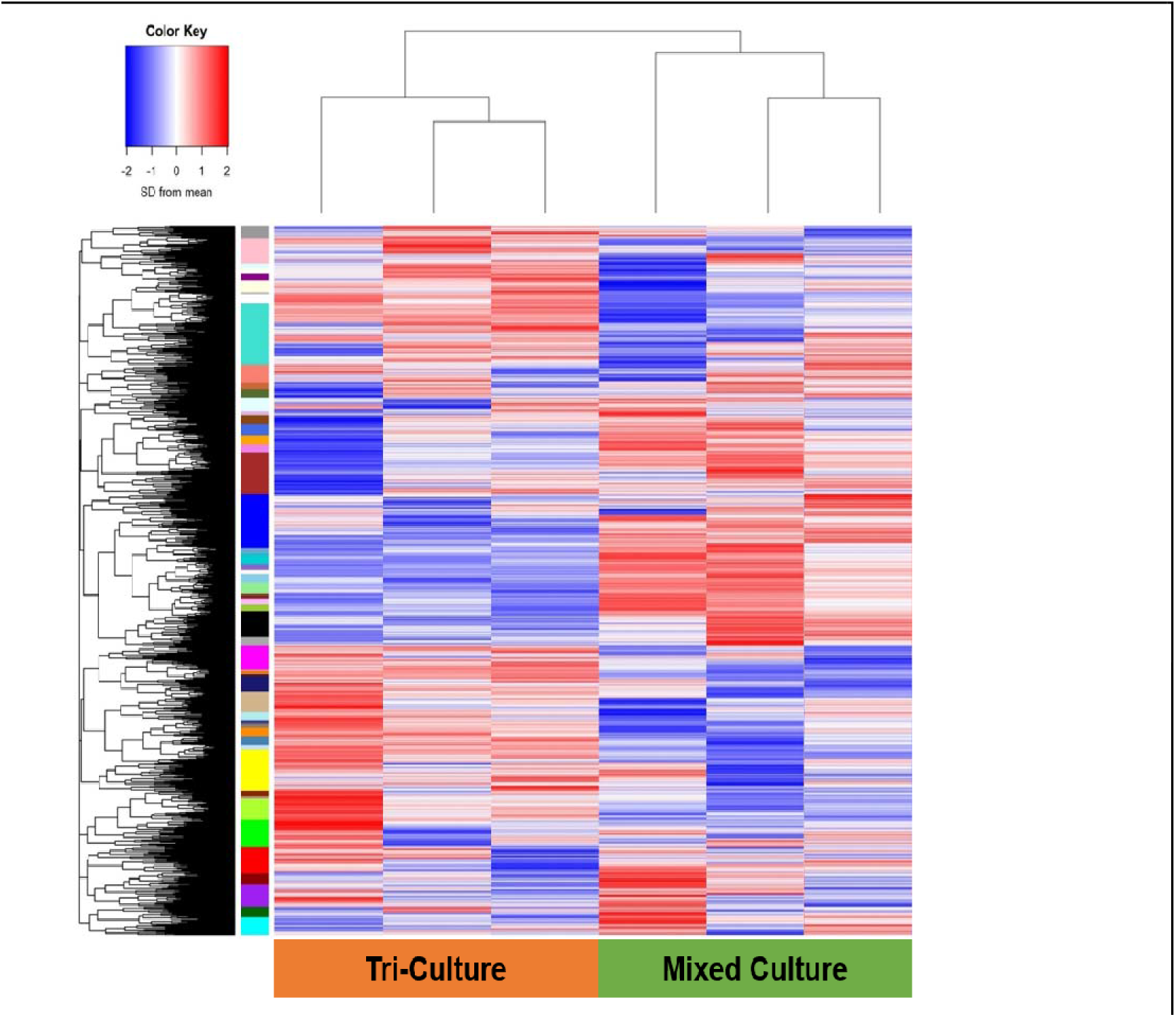
Expression pattern of genes detected during RNA-seq of *Tri-Culture* and *Mixed* samples. Color Key: SD from mean of log(Counts per Million) scaled such that values are centered at 0. N = 3.

**Figure 6.**
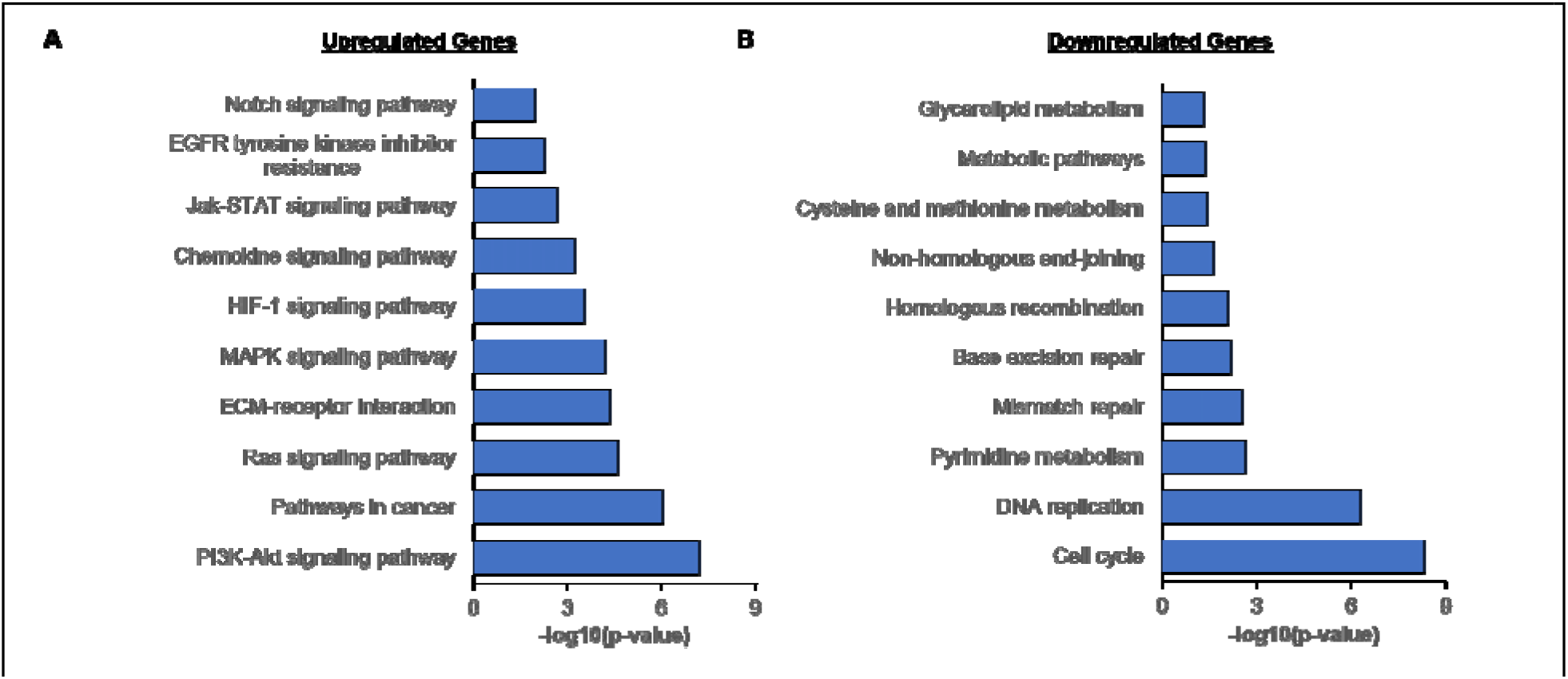
KEGG pathway analysis on statistically significant upregulated and downregulated genes. **(A)** Various pathways commonly dysregulated in cancer were over-represented amongst upregulated genes. **(B)** Pathways related to metabolism, DNA repair, and cell cycle were overrepresented amongst downregulated genes.

**Table 2.** Fraction of the cell population that is comprised of NHLFs in *Perivascular-only* and *Tri-Culture* hydrogels over the course of 14 days of culture. Data is presented as mean ± SD. N = 3.

| Day | CD90+ Percentage of Total Cell Population (%) |  |
| --- | --- | --- |
|  | Perivascular only | Tri-Culture |
| 3 | 57.72 ± 2.23 | 51.54 ± 1.14 |
| 7 | 52.25 ± 3.54 | 32.47 ± 0.58 |
| 14 | 34.62 ± 7.94 | 13.71 ± 0.81 |

**Table 3.** Gene Ontology (GO) terms that are over-represented amongst statistically significant upregulated genes.

| GO Term ID | P-Value | Term |
| --- | --- | --- |
| GO:0001944 | 3.62E-12 | vasculature development |
| GO:0030198 | 9.29E-11 | extracellular matrix organization |
| GO:0023052 | 1.88E-10 | signaling |
| GO:0030574 | 2.36E-09 | collagen catabolic process |
| GO:0042221 | 7.89E-09 | response to chemical |
| GO:0007155 | 3.29E-08 | cell adhesion |
| GO:0048584 | 4.16E-08 | positive regulation of response to stimulus |
| GO:2000145 | 9.58E-08 | regulation of cell motility |
| GO:0001525 | 4.98E-07 | angiogenesis |
| GO:0007166 | 5.85E-07 | cell surface receptor signaling pathway |
| GO:0009605 | 7.54E-07 | response to external stimulus |
| GO:0010646 | 2.39E-06 | regulation of cell communication |
| GO:0023051 | 3.75E-06 | regulation of signaling |
| GO:0023056 | 5.74E-06 | positive regulation of signaling |
| GO:0010647 | 6.03E-06 | positive regulation of cell communication |
| GO:0051272 | 0.0000127 | positive regulation of cellular component movement |
| GO:0046903 | 0.0000157 | secretion |
| GO:0022617 | 0.0000922 | extracellular matrix disassembly |
| GO:0023014 | 0.0001221 | signal transduction by protein phosphorylation |
| GO:0035690 | 0.000148 | cellular response to drug |
| GO:0001667 | 0.0001898 | ameboidal-type cell migration |
| GO:0009719 | 0.0002092 | response to endogenous stimulus |
| GO:0007167 | 0.0003346 | enzyme linked receptor protein signaling pathway |
| GO:0007169 | 0.0004987 | transmembrane receptor protein tyrosine kinase signaling pathway |
| GO:0090287 | 0.0005805 | regulation of cellular response to growth factor stimulus |
| GO:0051047 | 0.0006363 | positive regulation of secretion |
| GO:0071900 | 0.0008252 | regulation of protein serine/threonine kinase activity |
| GO:0040017 | 0.0008776 | positive regulation of locomotion |
| GO:0022407 | 0.0008782 | regulation of cell-cell adhesion |

**Table 4.** Gene Ontology (GO) terms that are over-represented amongst statistically significant downregulated genes.

| GO Term ID | P-Value | Term |
| --- | --- | --- |
| GO:0006270 | 1.04E-10 | DNA replication initiation |
| GO:0007049 | 2.46E-09 | cell cycle |
| GO:0051301 | 3.91E-08 | cell division |
| GO:0044770 | 0.00000125 | cell cycle phase transition |
| GO:0006139 | 0.00000313 | nucleobase-containing compound metabolic process |
| GO:1903047 | 0.00000838 | mitotic cell cycle process |
| GO:0000727 | 0.0000479 | double-strand break repair via break-induced replication |
| GO:0010564 | 0.0000485 | regulation of cell cycle process |
| GO:0006271 | 0.0000763 | DNA strand elongation involved in DNA replication |
| GO:0044237 | 0.0000823 | cellular metabolic process |
| GO:0000281 | 0.00010624 | mitotic cytokinesis |
| GO:1901988 | 0.00011111 | negative regulation of cell cycle phase transition |
| GO:0006268 | 0.00014277 | DNA unwinding involved in DNA replication |
| GO:0000082 | 0.00015003 | G1/S transition of mitotic cell cycle |
| GO:0000731 | 0.00019321 | DNA synthesis involved in DNA repair |
| GO:0007067 | 0.00021063 | mitotic nuclear division |
| GO:0051299 | 0.00037536 | centrosome separation |
| GO:0070987 | 0.00041795 | error-free translesion synthesis |
| GO:0006974 | 0.00042804 | cellular response to DNA damage stimulus |
| GO:0032467 | 0.00070774 | positive regulation of cytokinesis |
| GO:0046602 | 0.00078993 | regulation of mitotic centrosome separation |
| GO:0000070 | 0.00084012 | mitotic sister chromatid segregation |

### 3.4 Tri-culture and GBM-only hydrogels differentially respond to TMZ

The comparison of *Tri-culture* hydrogels to *Perivascular-only* and *GBM-only* hydrogels using RNA-seq revealed statistically significant upregulation of *MGMT* and downregulation of genes associated with DNA mismatch repair, which has been implicated in resistance to TMZ, the standard chemotherapy for GBM [62]. As such, we hypothesized that *Tri-culture* hydrogels would demonstrate increased resistance to TMZ compared to *GBM-only* hydrogels. Cell cultures were allowed to stabilize in GelMA and GelMA+HAMA hydrogels for seven days before treatment to allow the establishment of a perivascular niche-like environment within the *Tri-Culture* hydrogels. We then analyzed response to a 48 hour course of TMZ by assessing changes in metabolic activity and growth rate inhibition. Above a dosage of 250 μM TMZ, we observed that metabolic activity was lower in *GBM-only* hydrogels compared to *Tri-Culture* hydrogels (**Figure 7a**). We additionally used a cell-viability assay to measure growth rate inhibition in *GBM-only* and *Tri-Culture* hydrogels in response to TMZ treatment (**Figure 7b**). We did not observe growth rate (GR) inhibition in *Tri-Culture* hydrogels across the concentrations tested. In contrast, growth rate values decreased with increasing TMZ concentrations in *GBM-only* hydrogels. Using the Online GR Calculator (www.grcalculator.org/grcalculator), we estimated GR_50_ = 286 μM for *GBM-only* GelMA-only hydrogels and GR_50_ = 417 μM for *GBM-only* GelMA+HAMA hydrogels. In corroboration with trends observed with metabolic activity, we observed statistically significant reduced GR values for *GBM-only* cultures versus *Tri-Culture* environments containing a perivascular niche at dosages above 250 μM TMZ treatment (**Figure 7b, Supp. Figure 4**). Using imaging, we did not observe noticeable changes in the endothelial networks in the *Tri-Culture* platform before and after TMZ treatment (**Supp. Figure 5**).

**Figure 7.**
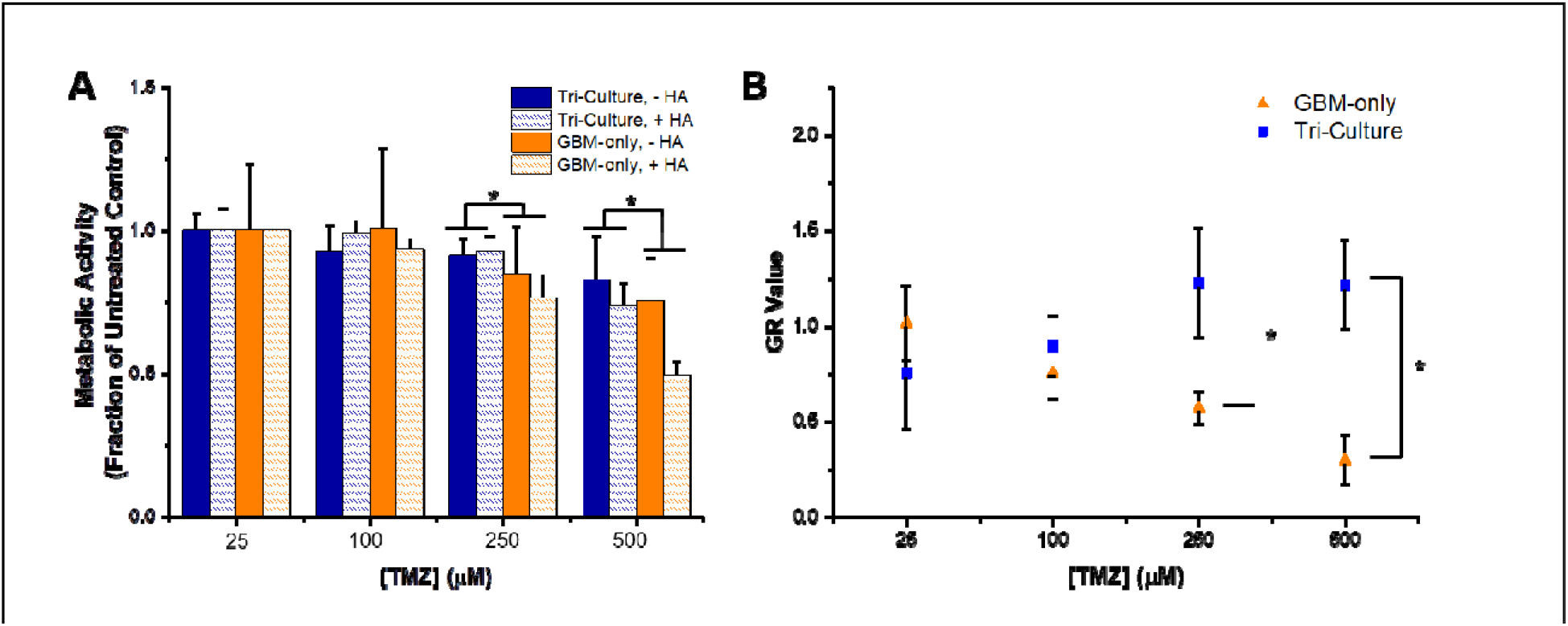
Comparison of response to TMZ in *GBM-only* and *Tri-Culture* hydrogels. **(A)** Metabolic activity (measured by MTT assay) was reduced in *GBM-only* cultures (higher response to TMZ) compared to *Tri-Culture* after treatment with 250 μM and 500 μM TMZ. *p<0.05 significant main effect of *Tri-culture* vs. *GBM-only* culture. The main effect only considers the culture platform by averaging across HA presence within a culture platform. **(B)** Growth rate (GR) inhibition occurred in *GBM-only* GelMA-only hydrogels with increasing concentrations of TMZ, but was not observed in *Tri-Culture* GelMA-only hydrogels. *p<0.05 between groups. Data is presented as mean ± SD. Two independent experiments were performed with three replicates each.

## 4. Discussion

GBM progression is believed to be linked to the interactions of cancer cells with endothelial cells and associated stromal cells within the perivascular niche. In particular, the spreading of GBM into healthy brain parenchyma is promoted by vessel co-option, regression, and angiogenic processes [11, 12, 15]. Previously, predominantly 2D culture or Transwell-based migration assays, as well as *in vivo* animal models, have been traditionally used to study interactions between GBM and endothelial cells [4, 63]. Limitations in these systems have lead to efforts in developing 3D *in vitro* models, and in particular there exists an opportunity to create platforms to study the impact of the perivascular niche environment on tumor progression.

We have previously demonstrated that endothelial cell networks can be sustained and tuned within gelatin hydrogels using a co-culture of endothelial cells and fibroblasts [43]. We also observed that including U87-MG tumor cells in the culture system lead to close association between tumor cells and the developing endothelial cell networks, followed by dissolution of these networks in a manner reminiscent of co-option. In this study, we quantified the kinetics of gene expression changes that accompanied the development and dissolution of the endothelial cell networks in response to culture with GBM cells over 14 days of culture *in vitro*. We were additionally interested in defining the degree to which the allowance for direct contact between heterogeneous cell types within the hydrogel culture altered the gene expression profile of the *in vitro* model compared to hydrogels containing each cell subpopulation individually. Based upon alterations in therapeutic response predicted by RNA-seq analysis, we examined whether the perivascular niche environment responded differently to temozolomide compared to hydrogels containing only tumor cells.

Vessel co-option and regression has been observed to precede sprouting angiogenesis at the tumor margins in GBM [12]. However, current *in vitro* platforms for studying endothelial and tumor cell interactions have not captured the kinetics of co-option and regression, and have spatially segregated the cell types to determine the ability of each cell type to induce migration or an angiogenic response [32, 34, 63–65]. By culturing endothelial, stromal, and tumor cells in direct contact within our platform, we observed behavior similar to angiogenesis followed by cooption and regression over a period of fourteen days. In native GBM, this sequence of events may occur at the tumor margins following vessel regression in the tumor core [12, 15]. The dynamics of *ANG2* and *VEGF* expression during co-option, regression, and angiogenesis have been described by *Holash et al*. and *Maisonpierre et al*. [12, 66]. Namely, in a rat model of GBM, *ANG2* expression with diffuse or absent *VEGF* expression was observed along with co-option and regression in the tumor core, while concurrent *ANG2* and *VEGF* expression was associated with angiogenesis and co-option at the tumor margins. In our platform, concurrent gene and protein expression of ANG2 and VEGF at Day 3 corroborated with the development of endothelial network structures. By Day 14, overall gene expression of *ANG2* had decreased, perhaps correlating with the reduced viability of endothelial cells as a result of regression. ANG2 protein expression by remaining endothelial cells along with diffuse VEGF expression also correlates with regressive behavior.

We investigated additional changes in gene expression accompanying endothelial network cooption and regression within the hydrogel platform. We measured gene expression related to receptor tyrosine kinases (RTKs), hyaluronic acid synthesis and interactions, angiogenesis, and ECM proteins and remodeling. RTKs encoded by *PDGFR, EGFR*, and *FGFR* drive signaling pathways that are commonly altered in GBM [6]. We observed decreases in gene expression with time in all RTKs studied except for *EGFR*. In particular, maximal expression of *VEGFR1, TIE2*, and *PDGFRβ* at Day 3 is tied to the initial angiogenic activity in developing endothelial networks, with *PDGFRβ/PDGFB* in particular have been implicated in the recruitment of pericytes to developing vessels [67] [68, 69]. Decreasing expression at subsequent time points correlates with regression, as these receptors are expressed on endothelial and stromal cells [70]. On the other hand, the constancy of *EGFR* expression reflects its enrichment in the tumor cell subpopulation [71]. With respect to genes related to hyaluronic acid synthesis, we observed higher *HAS1* and lower *HAS2* expression than that which was reported in PEG-based system containing only GBM tumor cells by *Wang et al*. [24]. *Wang et al*. additionally reported an increase in *HAS2* expression with time, while we observed an overall decreasing expression. In the presence of matrix immobilized hyaluronic acid within the hydrogels, we observed a biphasic trend in *HAS1* and *HAS2* expression, which mimics similar trends reported by *Pedron et al*. for *VEGF* and *HIF1α* [29]. Comparing with gelatin hydrogels containing only tumor cells, we noted similar patterns of expression for *VEGF* and *HIF1α* that were constant or trending upwards [29]. With the addition of HAMA, however, we did not observe biphasic trends reported previously, and instead noted increasing *VEGF* expression and *HIF1α* expression that trended downwards. The constant or decreasing *HIF1α* expression warrants further investigation, as increased *HIF-1* signaling has been implicated in triggering angiogenesis during and after regression [15]. With the lack of an angiogenic response after regression in our model, it may be possible that the early induction of *HIF1α* gives way to *HIF2α* expression for chronic hypoxia [72]. We additionally observed strong upregulation of *MMP9* during the culture period, consistent with previous studies reporting increased *MMP9* expression associated with heightened GBM aggressiveness in vascularized regions of the tumor [73, 74]. The observed increase in *MMP9* was greater in magnitude than that reported by *Wang et al*., and contrasted with the decline in expression reported by *Pedron et al*. [24, 29]. Finally, we noted upregulation of *TNC* during the culture period, which has been associated with tumor aggression and the perivascular environment [57, 75]. Overall, we have demonstrated that cellular diversity and the biomolecular composition of the biomaterial platform contributes to observed gene expression profiles. We acknowledge that gene expression is not necessarily directly correlated to the functional protein expression profile of the biomaterial; hence, future work is focused on determining protein expression dynamics during the development of the culture, particularly in response to therapeutics.

To further interrogate the impact of combined culture of perivascular niche cells in direct contact with tumor cells within the same biomaterial, we utilized RNA-seq to determine differentially expressed genes resulting from the combined culture of perivascular niche and tumor cells (*Tri-Culture*), compared to separate cultures of the cellular compartments. Analysis of differential gene expression revealed upregulation of processes relevant to the perivascular niche and GBM in the *Tri-culture* platform. For example, we observed upregulation of genes producing fibrillar matrix components prevalent in the basement membrane of vascular structures [76]. Additionally, genes involved in Notch signaling, which is implicated in cancer stem cell maintenance and angiogenesis within the perivascular niche, were upregulated [77, 78]. Direct culture of perivascular niche and tumor cells also resulted in enhanced signaling, notably in pathways identified by The Cancer Genome Atlas as being altered in a large percentage of GBM patients (PI3K/AKT, RTK/RAS) [6]. Specifically, we noted upregulation of *PDGFR, EGFR, FGFR1, and AKT3*, which are frequently amplified or activated in GBM. Additionally, within RB signaling, we observed downregulation of *CDKN2B* and upregulation of *CDK6*, which were inactivated in 47% and activated in 1% of The Cancer Genome Atlas dataset respectively [6]. These results suggest that while the *Tri-culture* platform does not recapitulate the native GBM PVN in its entirety, incorporating direct contact between multiple cell types within a biomaterial enhances select gene expression patterns relevant to GBM. In addition to enhanced signaling, upregulated genes were further correlated to the extracellular matrix, motility, and adhesion, while downregulated genes were related to cell cycle processes, reflective of the “go or grow” hypothesis. The relevance of this hypothesis has been demonstrated for GBM; *Mariani et al*. used a microarray analysis to demonstrate that genes related to cell cycle and proliferation were downregulated in migrating tumor cells [79]. Within the perivascular niche, *Farin et al*. demonstrated that while tumor cells migrated along vascular structures, they would stop movement in order to divide [11]. Tumor-associated endothelial cells have also been shown to be more migratory and less proliferative [80].

Furthermore, we compared the differential gene expression profile to results from other *in vivo* and *in vitro* models. *Khodarev et al*. used a microarray analysis to analyze gene expression in HUVECs co-cultured with U87-MG tumor cells, and reported upregulation in genes related to cell structure, motility, extracellular matrix, and receptors and their associated ligands [63]. *Bougnaud et al*. also used microarrays to compare gene expression profiles between various *in vivo* models of patient-derived xenografts, and showed upregulation of genes related to vascular development, ECM organization, and cell motion and adhesion from tumor cells isolated from an angiogenic xenograft [81]. We observe within our *in vitro* model that upregulated genes are correlated to similar biological processes. Interestingly, both *Khodarev et al*. and *Bougnaud et al*. noted upregulation of proliferation-related genes in endothelial cells, which contrasts with the overall downregulation of cell cycle processes observed in our model. This discrepancy may be explained by the fact that our model captures co-option and regression, and thus proliferative behavior by the endothelial cells is most likely observed during endothelial network formation in the beginning of the culture period. A deeper interrogation of temporal, cell type-specific gene and protein expression would enhance our understanding of this culture system, and is the subject of ongoing work. To interrogate genomic and secretome profiles of endothelial and tumor cell subpopulations individually, future efforts could employ techniques such as single-cell RNA-seq and heavy isotope labeling to study cell-cell interactions in GBM and other tumor types [71, 82, 83].

The ability for 3D *in vitro* models to model relevant aspects of GBM behavior presents opportunities for such platforms to be used to investigate drug response in an efficient manner and potentially facilitate drug discovery. The current standard chemotherapy for GBM is temozolomide (TMZ), and *MGMT* expression as well as the dysregulation of the DNA mismatch repair (MMR) pathway are two mechanisms implicated in TMZ resistance [62]. Notably, *MSH6* was inactivated in 4% of the TCGA dataset [6]. We confirmed heightened resistance to TMZ in the *Tri-culture* compared to the *GBM-only* model using metabolic and cell viability assays. The differential response to TMZ observed between the two models corroborates with the observed upregulation of *MGMT* and downregulation of MMR-related genes in the *Tri-culture*, as established through our RNA-seq analysis. The perivascular niche has been associated with therapeutic resistance, partially due to its ability to maintain a resistant cancer stem cell population [18], however in this study the GBM cell line we employed does not contain a measurable GBM stem cell population (not shown). Studies have suggested that endothelial cells are resistant to TMZ, which would additionally contribute to resistance [84, 85]. Taken together, these results demonstrate that the tri-culture model recapitulates heightened therapeutic resistance associated with the perivascular niche, and represents a promising opportunity for future studies that deduce signaling pathways contributing to resistance and propose potential inhibitors to improve therapeutic response.

Based on this work, we acknowledge limitations that provide opportunities for future work to improve and understand the recapitulation of the perivascular niche *in vitro*. First, we acknowledge that the process of hydrogel dissociation to retrieve cells for flow cytometry analysis can potentially alter surface receptor and signaling profiles. Thus, we are interested in utilizing biomaterial platforms with rapidly reversible crosslinking to enable facile collection of cells for post-culture analysis [86]. Moreover, we acknowledge that the physiological relevance of this platform can be further enhanced by utilizing a variety of primary tumor specimens in addition to brain-specific perivascular cells. The transition from GBM cell lines to patient-derived cells provides an additional opportunity to study the behavior of cancer stem cells, their interactions with artificially created perivascular niches within a hydrogel environment, and their role in therapeutic response.

## 5. Conclusions

The perivascular niche in GBM has been implicated in contributing to tumor invasion and therapeutic resistance. Incorporating elements of the perivascular niche into 3D *in vitro* platforms is therefore instrumental in improving the usefulness of this culture system for understanding tumor biology and accurately predicting therapeutic outcome. We report that the inclusion of endothelial cells and fibroblasts in co-culture with U87-MG cells within GelMA hydrogels produces a gene expression profile that allows for interrogation of biological processes and therapeutic response relevant to GBM. Upregulation of genes related to motility and signaling alongside downregulation of genes related to cell cycle point towards a model that supports the “go or grow” phenotype. Temporal changes in gene expression additionally supported observations of endothelial network formation, co-option, and regression during the culture period. Finally, we observed enhanced resistance to the alkylating agent TMZ from the synergistic tri-culture in GelMA hydrogels as opposed to the culture of tumor cells alone.

Overall, this platform will be useful for studies that examine mechanisms governing associations between GBM and endothelial cells that eventually lead to invasion, as well as those that contribute to therapeutic resistance within the perivascular niche.

## Acknowledgements

The authors would like to acknowledge the members of the Roy J. Carver Biotechnology Center at the University of Illinois Urbana-Champaign for their advice and assistance with experiments for this manuscript. Specifically, the authors would like to thank Drs. Barbara Pilas and Angela Kouris in the Flow Cytometry Facility, Dr. Mark Band and Tatsiana Akraiko in the Functional Genomics Unit, Dr. Alvaro Hernandez and Chris Wright in the High-Throughput Sequencing and Genotyping Unit, and Drs. Chris Fields and Jenny Zadeh in the High-Performance Biological Computing Group. The authors also gratefully acknowledge the Computer and Networking Resource Group at the Carl R. Woese Institute for Genomic Biology at the University of Illinois at Urbana-Champaign for assistance in setting up resources for high-performance computing for RNA-seq analysis. Research reported in this publication was also supported by the National Cancer Institute of the National Institutes of Health under Award Number R01 CA197488. The content is solely the responsibility of the authors and does not necessarily represent the official views of the NIH. We are grateful for the funding for this study provided by the NSF Graduate Research Fellowship DGE-1144245 (MTN). The authors are also grateful for additional funding provided by the Department of Chemical & Biomolecular Engineering and the Carl R. Woese Institute for Genomic Biology at the University of Illinois at Urbana-Champaign.

## Data availability

All data analyzed during this study are included in this published article (and its supplementary information files). Raw RNA-seq data can be viewed on the NCBI GEO repository using accession number GSE111231. Other raw data required to reproduce these findings are available from the corresponding author on reasonable request.

---

**Supplementary Table 1**

Statistically significantly upregulated genes in *Tri-culture* RNA samples compared to *Mixed* RNA samples. Statistical significance is determined by FDR-adjusted p-value<0.05. N = 3.

**Supplementary Table 2**

Statistically significantly downregulated genes in *Tri-culture* RNA samples compared to *Mixed* RNA samples. Statistical significance is determined by FDR-adjusted p-value<0.05. N = 3.

**Supplementary Table 3**

Gene Ontology (GO) terms related to biological processes that are over-represented amongst statistically significant upregulated genes within *Tri-culture* RNA samples compared to *Mixed* RNA samples. p<0.05 for statistically significantly over-represented terms.

**Supplementary Table 4**

Gene Ontology (GO) terms related to biological processes that are over-represented amongst statistically significant downregulated genes within *Tri-culture* RNA samples compared to *Mixed* RNA samples. p<0.05 for statistically significant over-represented terms.

**Supplementary Table 5**

KEGG pathways that are over-represented amongst statistically significant upregulated genes within *Tri-culture* RNA samples compared to *Mixed* RNA samples. p<0.05 for statistically significant over-represented pathways.

**Supplementary Table 6**

KEGG pathways that are over-represented amongst statistically significant downregulated genes within *Tri-culture* RNA samples compared to *Mixed* RNA samples. p<0.05 for statistically significant over-represented pathways.

